# Epipelic biofilms respond early to the impacts of urbanization in lowland streams

**DOI:** 10.1101/2020.06.22.164715

**Authors:** Joaquín Cochero, María Mercedes Nicolosi Gelis

## Abstract

Multiple structural and functional endpoints of stream biofilms are employed by water quality monitoring programs to detect both direct and indirect environmental impacts. Since multiple co-occurring stressors influence biofilm development, active biomonitoring (translocation experiments) could provide a useful monitoring tool that reflects the overall water quality of the urbanized sites. The aim of this research was to study the short-term responses of epipelic biofilms caused by their translocation to more polluted reaches in lowland urban streams. Fluvial sediment was translocated in three streams that run through urban areas following an urbanization gradient. The epipelic biofilms in the sediment were sampled to identify any fast occurring changes in their algal and bacterial biomasses, in their respiration and oxygen consumption. The results show that structural changes in the biofilm, such as an increased bacterial density and chlorophyll-a concentration, were measurable after two days of exposure to sites with impaired water quality. These immediate changes in the structure of the biofilm indicate that they are sensitive endpoints that can be employed in fast and inexpensive biomonitoring programs in urbanized streams.

## 1. INTRODUCTION

Fluvial biofilms (Lock 1981) have been used to monitor physical and chemical water quality changes since the short life cycle of the microorganisms and the trophic interactions between the microbiota allow for the detection of both short and long-term impacts. These communities are among the first to react to environmental stressors, including nutrient enrichments (Guasch et al. 1995; Rier and Stevenson 2002; Olapade and Leff 2005), heavy metals (Ramelow et al. 1987; Medley and Clements 1998; Giorgi and Malacalza 2002; Sierra & Gómez 2010), pesticides and herbicides (Dorigo et al. 2007; Tlili et al. 2008; Ricart et al. 2010) and pharmaceutical components(Proia 2012).

The process of urbanization affecting streams and rivers is linked to the effects of multiple simultaneous stressors, the “urban stream syndrome” (Walsh et al. 2005), which confounds the ability to make inferences about the relative effect of each stressor on the biota (Paul and Meyer 2001, Walsh et al. 2005b). Hence, multiple structural and functional endpoints from the biofilm are employed in water quality monitoring programs of both direct and indirect environmental impacts (Proia et al. 2012). For instance, water quality detriments can lead to an increase in algal biomass and algal pigments (Tien et al. 2009), in bacterial growth (Leff 2000; Tien et al. 2009), in metabolic activity (Guasch et al. 1995; Garnier and Billen 2007), and are usually linked to a modification in the specific composition (Rimet et al. 2009; Delgado et al. 2012, Morin et al. 2016).

Even in environments with high availability of resources, such as nutrient rich streams (Feijoó et al. 1999; Bauer et al. 2002; Licursi et al. 2016), biofilms have shown to respond to slight variations in water quality related to the effects of urbanization, by developing a greater biomass with a higher metabolic activity (Vilches 2012; Cochero et al. 2013, 2018). Translocation assays (also known as “active biomonitoring”) employing biofilms could represent a sensitive monitoring tool that reflects the overall water quality of the sites even when multiple co-occurring stressors are present. These biomonitoring techniques consist of the transfer of organisms from one place to another to quantifying their biochemical, physiological and/or organismal responses for the purpose of water quality monitoring (De Kock and Kramer 1994). The use of translocation approaches employing biofilms have been used successfully to describe the effects of various environmental and anthropogenic stressors, yet only a few related to organic pollution in urban streams (Ivorra et al. 1999; Sierra and Gómez 2010; Tlili et al. 2011; Proia et al. 2013b, a; Nicolosi Gelis et al. 2020), and the large majority in nutrient-depleted fluvial systems with epilithic substrates.

The aim of this research was to study if there were short-term responses (up to 72hs) measurable in the epipelic biofilm when translocated from peri-urban sites to more urbanized reaches in lowland streams. We hypothesized that, since the structure and metabolism of biofilms are rapidly affected by the physical-chemical characteristics of the water, they would reflect the environmental changes quickly. The increase in organic matter and nutrients would therefore be reflected as increments in bacterial density, chlorophyll*-α* content and in the metabolic activity of the translocated biofilms.

## 2. MATERIALS AND METHODS

### 2.1 Sites selection and physical-chemical data

To conduct the translocation assays, three streams (“Don Carlos” stream, hereafter “Stream 1”; “El Gato” stream, hereafter “Stream 2”; “Rodriguez” stream, hereafter “Stream 3”) that run through urbanized areas near the city of La Plata (Argentina) were selected *a priori* from previous research considering their degree of exposure to urbanization (Cochero et al. 2016, 2018). Briefly, their degree of urbanization was measured by calculating the percentage of impervious surface in the watershed related to urban activities, by obtaining land-use maps (Instituto Geográfico Nacional, www.ign.gob.ar).

Two sites were selected from each stream: a “Peri-urban” site, located upstream on the outskirts of the city and exposed to mild agricultural activities, and an “Urban” site, located downstream, well within the urbanized area of La Plata city (site coordinates in Figure 1). All sites had similar hydrological characteristics, low velocity (<0.02 m s^-1^), similar turbidity (90±32NTU) and they lack riparian vegetation and macrophytes.

**Figure 1.**
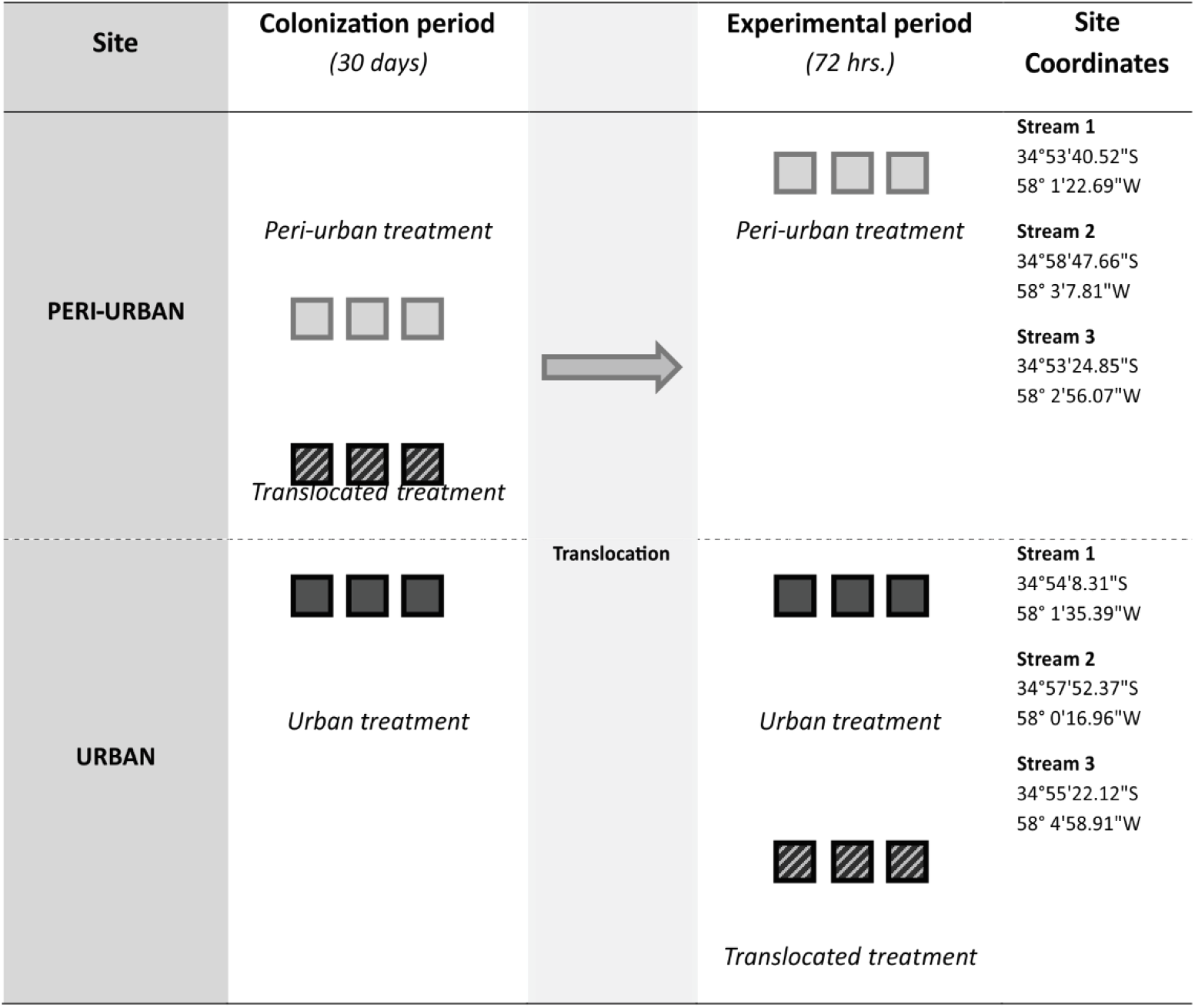
Experimental design used involving the translocation of germination trays with sediment from a peri-urban site to an urban site within streams. The Peri-urban treatment trays (light gray squares) were colonized in the Peri-urban sites and kept there as controls; the Urban treatment trays (dark gray squares) were colonized in the Urban sites and kept there as controls; and the Translocated treatment trays (striped squares) were colonized in the Peri-urban sites and transferred to the Urban sites for the experimental period.

Dissolved Oxygen (DO, mgL^-1^), temperature (°C), conductivity (μS cm^-1^), pH and turbidity (NTU) were measured at each site using a multiparametric sensor (Horiba U50). Water samples (500 ml) were collected in the Peri-urban and Urban sites at all sampling times in order to measure nutrients (ammonium, nitrites, nitrates, soluble reactive phosphorous), biochemical oxygen demand (BOD_5_, mgL^-1^) and chemical oxygen demand (COD, mgL^-1^) samples, which were analyzed according to standardized spectrophotometric methods (APHA/AWWA/WEF, 2005).

### 2.2 Experimental setup

Nine germination trays (200 wells, 2.5 diameter x 3cm deep each well) were filled with stream sediment and attached to the streambed ensuring light penetration and left to be colonized for 30 days at each stream. Within each stream, six of the trays were placed in the upstream (Peri-urban) site and the other three trays were placed in the downstream (Urban) site. After the colonization period, three of the trays from the Peri-urban site were transferred in a humid environment to the Urban site and attached to the streambed. Therefore, three treatments were established: control trays for the Peri-urban sites (named “Peri-urban treatment”), control trays for the downstream sites (named “Urban treatment”) and the trays that were transferred from the Peri-urban sites to the Urban sites (named “Translocated treatment”) (Figure 1).

Every 24hs three epipelic biofilm subsamples were collected at random from each tray by pipetting the surface layer (0.5cm) of the sediment (Gómez et al. 2011) and transported to the lab in coolants for analysis. Biological endpoints were measured in all three subsamples and their values were averaged for the statistical analyses; therefore, each tray acted as an experimental unit. Results obtained from the biological endpoints (Chlorophyll-*a*, bacterial density, ETS activity, O_2_ consumption) were referred to cm^2^ of sampled sediment by a granulometric analysis (Folk 1980). The experimental period lasted 72hrs.

#### Bacterial density

Samples were stored in sterile glass vials with 2% v/v formalin. Bacterial density was estimated after sonication (three 2 min cycles) and appropriate dilution (1:100 to 1:400) of the samples. Diluted samples were stained for 10 min with DAPI (4’, 6-diamidino-2-phenyilindole) to a final concentration of 1 μg mL^-1^ (Porter and Feig 1980), and filtered through a 0.2 μm black polycarbonate filter (GE Osmonics). Bacteria were then counted using an epifluorescence microscope (Olympus BX-50) under 1000× magnification and with an Olympus Q-Color5 imaging system. Twenty fields per replicate were counted for a total of 400 to 800 organisms.

### 2.3 Chlorophyll-a

Chlorophyll*-a* content was measured by spectrophotometric methods. Samples were first sonicated for three 2-minute cycles in a Cleanson CS1106 sonicator and filtered through Sartorious GF/C filters. Chlorophyll*-a* was extracted with 90% acetone for 12 h and the supernatant was read in a Labomed UV-VIS Auto 2602 spectrophotometer. Chlorophyll-*a* concentration (mg cm^-2^) was calculated according to (Strickland and Parsons 1972).

### 2.4 Activity of the electron transport system (ETS)

The *ETS activity* represents a measure of the overall respiration of the biofilm, and was assayed by measuring the reduction of the electron transport acceptor INT (2-3 tetrazolium chloride) into INT-formazan (iodonitrotetrazolium formazan) (Blenkinsopp and Lock 1990). Samples were incubated in 0.02% INT solution *(Sigma-Aldrich)* for 12 hours in the dark. INT-formazan was then extracted with cold methanol for a minimum of 1 h at 4°C in the dark followed by sonication (Cleanson CS1106). Absorbance at 480 nm was measured in a Labomed UV-VIS Auto 2602 spectrophotometer and converted to ETS activity values (μg INT-formazan cm^-2^ h^-1^) using a and a stock solution of 30 mg mL^-1^ INT-formazan in methanol.

### 2.5 Additional oxygen consumption of the sediment

The *additional oxygen consumption* method (Knopp 1968) uses a substrate to stimulate bacterial growth (peptone) in the samples, and assumes that if the bacterial activity is normal, the respiration associated with the reduction of the additional substrate leads to an increased oxygen consumption. This can be measured as a greater oxygen depletion at the end of an incubation period. If bacteria are being inhibited by a stressor, oxygen consumption ceases or lowers significantly. The additional oxygen consumption (or “ZZ”, for its acronym in German) therefore provides a fast and comparative measurement of the overall bacterial respiratory activity. In these assays, samples were incubated for 24hs after a peptone (50 mg L^-1^) spike was added, and the dissolved oxygen consumed was compared to control samples without peptone. Results are expressed as the amount of oxygen consumed by the biofilm per hour (mg O_2_ cm^-2^ h^-1^).

### 2.6 Statistical analysis

Differences in physical-chemical variables between sites were analyzed considering the sites as replicates depending on their degree of urban impact (Peri-urban n=3; Urban n=3). A two factor Multivariate Analysis of Variance (MANOVA; *Time:* 0, 24hs, 48hs, 72hs; *Site:* Peri-urban, Urban), following the model [X = μ + Site + Time + Time * Site + Residuals]. The biological variables from the biofilms were also analyzed by a two factor MANOVA (*Time*: 0, 24hs, 48hs, 72hs; *Treatment:* Peri-urban, Urban, Translocated), following the model [X = μ + Treatment + Time + Time * Treatment + Residuals].

Values were first transformed to log(x+1) to ensure normality, which was previously assessed by the Shapiro-Wilk test (Shapiro and Wilk 1965); homogeneity of variance was tested by Cochran’s C test (Cochran 1951). For each biological variable, three subsamples from each tray were collected, and their mean value was used for the statistical analyses.

The Student-Newman-Keuls (SNK) post-hoc test was used, and partial eta^2^ (η^2^) was computed as a measure of the effect size.

## 3. RESULTS

### 3.1 Physical-chemical data

Results from the multivariate analyses show significant differences between Peri-urban and Urban sites (Site factor: F=10.19, p<0.05; Partial μ^2^=0.96), regardless of the sampling time (Time factor: F=2.03, p=0.07, partial μ^2^=0.82).

Univariate analyses for each physical-chemical variable show that, when compared to the Peri-urban sites, the Urban sites had significantly higher concentrations of DIN (+80.7%), SRP (+234%), BOD_5_ (+97.8%), and higher conductivity values (+62.9%) (Table 1, Figure 2). The effect size measure shows that the differences in physical-chemical variables between sites is intermediate (Partial 0.75 < μ^2^ ≥ 0.25).

**Figure 2.**
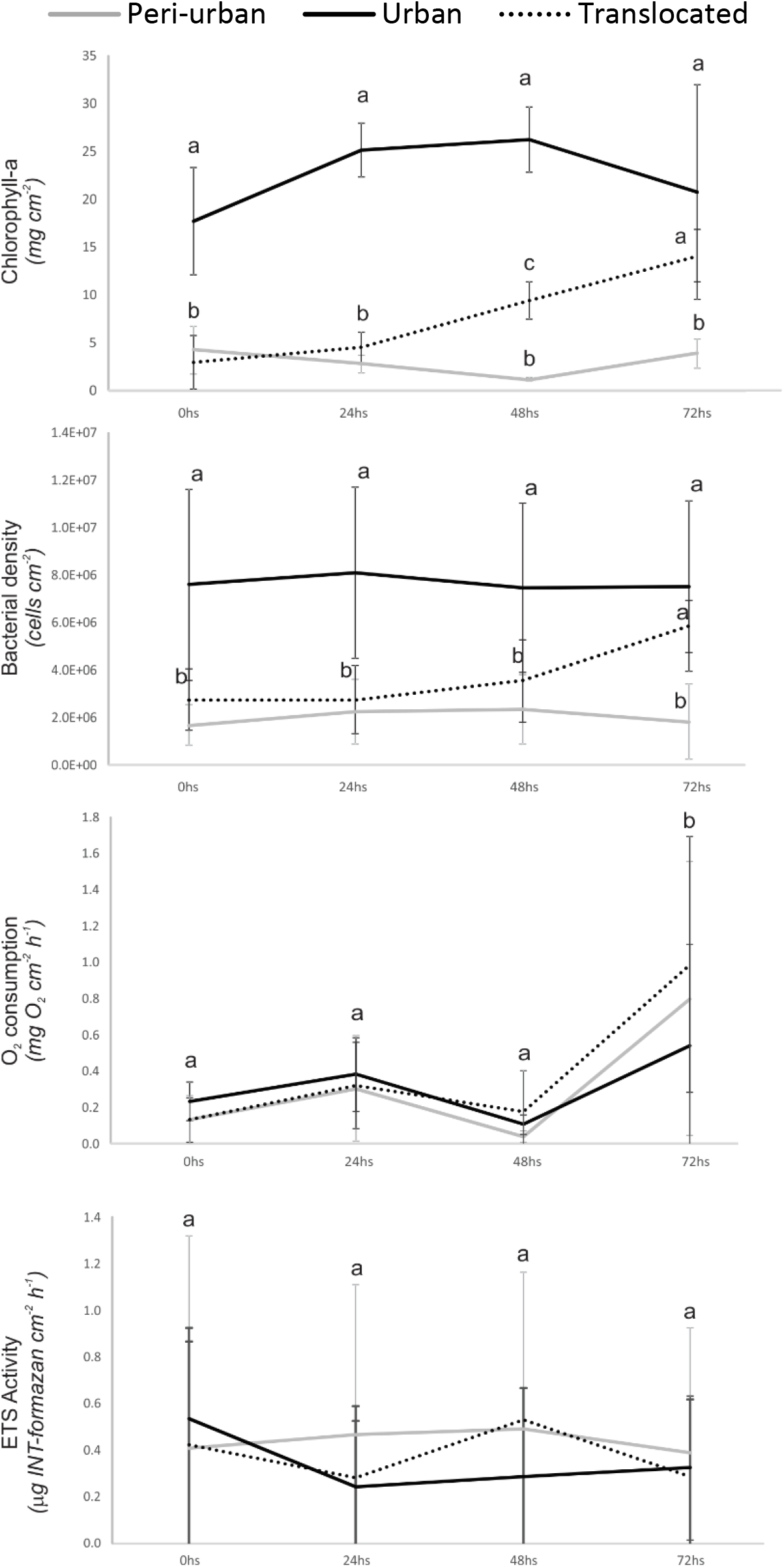
Mean (±SD) chlorophyll-*a* (*mg cm*^-2^), bacterial density (*cells cm*^-2^) ETS activity (*μg INT-formazan cm*^-2^ *hr*^-1^) and O_2_ consumption (*mg O*_2_ *cm*^-2^ *hr*-^1^) during the experiment in the Peri-urban (black solid line), Urban (gray solid lines) and Translocated (black dotted line) treatments. Letters represent groups obtained from the post-hoc statistical analyses at each sampling time comparting between treatments.

**Table 1.** Mean (± SD) values obtained for the physical-chemical variables in the Peri-urban and Urban sites, along with their mean increment between sites (%), the significance values from the MANOVA (Site Factor) pairwise comparisons and a measure of the effect size (partial μ^2^).

| Variable | Peri-urban | Urban | Increment (%) | Significance ($p$-value) | Effect size (Partial $\mu^2$) |
| --- | --- | --- | --- | --- | --- |
| **DIN** (mg L<sup>-1</sup>) | 1.1 ($\pm 1.01$) | 1.99 ($\pm 0.65$) | 80.7 | **0.031** | 0.259 |
| **SRP** (mg L<sup>-1</sup>) | 0.41 ($\pm 0.18$) | 1.37 ($\pm 1.31$) | 234.0 | **0.046** | 0.226 |
| **BOD<sub>5</sub>** (mg L<sup>-1</sup>) | 7.5 ($\pm 4.12$) | 14.83 ($\pm 5.97$) | 97.8 | **0.005** | 0.400 |
| **COD** (mg L<sup>-1</sup>) | 19.36 ($\pm 10.22$) | 27.58 ($\pm 10.77$) | 42.5 | 0.074 | 0.186 |
| **Temperature** (°C) | 11.5 ($\pm 2.6$) | 13.4 ($\pm 2.5$) | 15.9 | 0.094 | 0.165 |
| **pH** | 7.57 ($\pm 0.49$) | 7.81 ($\pm 0.33$) | 3.2 | 0.329 | 0.060 |
| **Turbidity** (NTU) | 91.2 ($\pm 99.7$) | 62.74 ($\pm 59.3$) | -31.2 | 0.465 | 0.034 |
| **Conductivity** ($\mu\text{S cm}^{-1}$) | 479.5 ($\pm 270.3$) | 781.1 ($\pm 230.3$) | 62.9 | **0.014** | 0.321 |
| **DO** (mg L<sup>-1</sup>) | 5.61 ($\pm 2.14$) | 5.91 ($\pm 2.65$) | 5.4 | 0.710 | 0.009 |

### 3.2 Biofilms

Results from the multivariate analyses show a significant interaction between the Treatment (Peri-urban, Urban, Translocated) and Time (0, 24hs, 48hs, 72hs) factors (MANOVA Treatment*Time F=1.751, p<0.05; Partial μ^2^=0.13).

Chlorophyll-*a* concentration in the biofilms was higher in the Urban sites than in the Periurban sites (F=85.46; p<0.05). At the time of the translocation (time 0hs), the chlorophyll-*a* concentration in the Translocated biofilm was similar to the concentration in the Periurban biofilm (p>0.05). After 48hs the chlorophyll*-a* concentration in the Translocated biofilm was significantly different from that at the Peri-urban biofilm, and after 72hs of exposure the algal biomass in the Translocated treatment was similar to the Urban treatment (pairwise comparisons p<0.05; Figure 2).

The variations in bacterial density also followed a similar pattern to that of the algal biomass (Figure 2). At time 0hs bacterial density was higher in the Urban biofilm than both in the Peri-urban and Translocated biofilms. After 72hs from the translocation, the bacterial density in the biofilm had increased to resemble the downstream biofilm (pairwise comparisons p<0.05, Figure 2).

Both ETS activity and O_2_ consumption were highly variable during the experiment (Figure 2), and there were no significant differences between treatments at any date (Table 2). There was, however, an increase in O_2_ consumption in the final sampling time in all treatments (F=17.8; p<0.05).

**Table 2.** MANOVA between-subjects results and effect size measure (μ^2^) for the biological variables measured during the experiment (*p <0.05).

|  |  | Treatment | Time | Treatment * Time |
| --- | --- | --- | --- | --- |
| **Bacterial density** | F | 52.04 | 0.89 | 1.35 |
|  | p | **<0.01\*** | 0.45 | 0.24 |
| | $\mu^2$ | 0.52 | 0.03 | 0.08 |
| **Chlorophyll-*a*** | F | 85.46 | 4.70 | 4.36 |
|  | p | **<0.01\*** | **<0.01\*** | **<0.01\*** |
| | $\mu^2$ | 0.71 | 0.17 | 0.27 |
| **ETS activity** | F | 0.29 | 0.45 | 0.34 |
|  | p | 0.75 | 0.72 | 0.91 |
| | $\mu^2$ | 0.01 | 0.01 | 0.02 |
| **O<sub>2</sub> consumption** | F | 0.66 | 17.79 | 1.09 |
|  | p | 0.52 | **<0.01\*** | 0.37 |
| | $\mu^2$ | 0.01 | 0.36 | 0.06 |

## 4. DISCUSSION

The results presented here show that even after a short exposure, the translocation assay can result in measureable endpoints in the biofilm that respond to the negative impacts of urbanization (*i.e*. increased nutrients and organic matter concentrations)

Nutrient availability is one of the most important factors in biofilm development, even in nutrient-rich streams with high contents of organic matter (Carr et al. 2005; Cochero et al. 2013). Since Pampean streams are rich in phosphorous and nitrogen, it could be possible that the microbial communities are carbon-limited. The transfer of biofilms to a site with higher concentrations of BOD_5_ and COD would therefore translate to higher biomasses. In this study, after only 48hs of exposure the transferred biofilm exhibited a greater biomass development of both the bacterial and algal communities. Bacterial density increased significantly after 72hs of exposure, while algal biomass (i.e. chlorophyll-a) started spiking after 48hs of exposure. The short delay between algal and bacterial increases in the transferred biofilm could indicate that this new source of carbon in the more urbanized sites does not represent a high quality source for the bacterial assemblage, and bacteria would therefore incorporate carbon exudates from the algal community. This is consistent with previous results obtained in Pampean streams, where the carbon-limited bacterial community in the sediment was tightly linked to the accrual of the phytobenthos (Cochero et al. 2013).

Metabolic variables usually react sooner to environmental impacts than structural characteristics (Serra et al. 2009; Guasch et al. 2010; Cornet 2012; Proia et al. 2013a), and the effects are generally more evident when the biofilm from a less polluted site is exposed to heavier pollution than the other way around (Sierra and Gómez 2010; Proia et al. 2013b). For instance, the primary production of the biofilm decreased when transferred from a less polluted site to a more polluted site in a nutrient-rich stream before any changes were measurable in algal biomass (Sierra and Gómez 2010). Our results show that the respiration measured both as the activity of the electron-transfer system and as sediment oxygen consumption, were highly variable endpoints throughout the experiment, not clearly allowing the discrimination between treatments. This metabolic variability could be related to the stress to which the biofilm is subject to due to the translocation process itself, rather than a natural variation in the communities, and should be taken into account in future monitoring protocols. In long term experiments, metabolic variables have shown to stabilize (Sabater et al. 2011), linking streams exposed to urban impacts to a higher heterotrophic activity in the sediment (Bukaveckas et al. 2002; Artigas et al. 2013; Aristi et al. 2015). However, respiration endpoints from the sediment should be analyzed with caution for short-term translocation experiments aimed to recognize fast changes in water quality conditions.

The use of translocation experiments has been increasingly used for communities such as diatoms (Ivorra et al. 1999; Tolcach and Gómez 2002; Sierra and Gómez 2010; Nicolosi Gelis et al 2020), as they can represent a powerful approach for the biomonitoring of watersheds, providing a suitable design for multiple stressor assessment. Similar translocation assays have been conducted before in the field, although usually using artificial substrates such as glass (Ivorra et al. 1999; Arini et al. 2012; Proia et al. 2013b, a), while almost no research has been conducted in lowland streams where the predominant substrate are sediments with large contents of silts and clay. By using germination trays with stream sediment, we ensure that the translocated assemblage is similar to that found in the stream, therefore providing more realistic results than when introducing an artificial substrate such as glass or plastic into the stream. This represents a clear advantage when using translocation experiments to further the knowledge of how sediment biofilms react to changes in environmental conditions.

## 5. CONCLUSIONS

This study highlighted the sensitivity of fluvial biofilms to respond to change in a short exposure period in urbanized streams, even when the change in water quality is small or moderate. The translocation assay was an effective strategy to measure how some structural parameters from the biofilm represented the gradient from peri-urban to urban sites. The metabolic endpoints used in these assays were highly variable and were difficult to relate to the environmental change, but the algal and bacterial biomasses were stable parameters to follow these changes in environmental conditions. Further uses of these translocation experiments could offer a more detailed idea of how specific physical and chemical conditions affect fluvial biofilms, how they recover from environmental impacts and which endpoints can be further explored for fast and inexpensive biomonitoring purposes.

## ACKNOWLEDGEMENTS

This work was funded by the Consejo Nacional de Investigaciones Científicas y Técnicas of Argentina [Grant number PIP-173].

